# Free meals during breeding: increased resource access does not benefit Arctic-nesting shorebirds

**DOI:** 10.1101/2022.05.30.494000

**Authors:** Laurent Montagano, Nicolas Lecomte

## Abstract

Arctic ecosystems are facing intensifying impacts of climate change, notably an increase in air temperature that can boost local productivity. Yet whether such resource surge can benefit declining migratory birds is unclear. Here we experimentally increased prey abundance and measure its effect on the body condition, nesting patterns, and nesting success of the white-rumped sandpiper (*Calidris fuscicollis*). We captured females at the beginning and at the end of their incubation to assess body condition and we installed small temperature probes inside their nests to measure their incubation recess. To estimate nest survival, we regularly monitored experimental and control nests during two consecutive summers (2016–2017). For both experimental years with contrasting nest success (73% vs. 21%), we found no evidence of an effect of our supplementation experiment mimicking an increased abundance of arthropods in the Canadian Arctic. This suggests that *in situ* resources are not limiting during incubation. Breeding strategies and success in shorebirds seem to be driven by inter-individual traits related to body condition upon the initiation of incubation.

## Introduction

The migration of Arctic-breeding birds often entails high energetic costs related to travelling great distances (Klaassen 1996). Upon arrival on their arctic breeding grounds, income breeders (Drent and Daan 1980, Morrison and Hobson 2004) like sandpipers must regain fat reserves and reconstitute their reproductive organs as quickly as possible. Unlike some other birds, they must also feed during incubation as well. On the opposite end of the spectrum, purely capital breeders such as eiders rely exclusively on their energy stores accumulated prior to arriving on the breeding grounds (Drent & Daan 1980; Morrison & Hobson 2004). However, for most other capital and income breeders, the quantities of both body stores upon arrival and *in situ* resources are important. Thus, carry-over effects influencing the physical shape of birds when they arrive in the Arctic such as migration (Legagneux et al. 2012, 2016) and wintering (Marra et al. 1998) combined with feeding strategies and the availability of resources on the breeding ground should be critical components influencing incubation and nesting success of Arctic-breeding birds.

Incubation is a complex period as one or both parents must maintain the temperature of their eggs above a threshold, deter nest predators, and feed. Generally, in order for embryos to develop correctly, nest temperatures should not drop below a temperature threshold of 35 °C (White and Kinney 1974). In High Arctic environments where average air temperatures usually range from −10 to 10 °C in the summer, this means that birds often cannot leave their nest unattended for long periods of time (Cantar and Montgomerie 1985, Olson et al. 2006). However, income breeders also regularly need to feed in order to maintain sufficient body stores reserves (Cresswell 2004, Reneerkens et al. 2011, Bulla et al. 2016) and compensate for high daily energy expenditure (Piersma et al. 2003). Cold Arctic temperatures, wind, and rain intensify energetic demands and have been shown to modulate the frequency of foraging recesses of income breeders (Cantar and Montgomerie 1985, Tulp and Schekkerman 2006, Lecomte et al. 2009). Despite lower predation pressure at higher latitudes (McKinnon et al. 2010), avian and terrestrial nest predators can cause depredation to be the leading cause of nest failure (Bêty et al. 2001, Weiser et al. 2017). Parents can limit these impacts by actively deterring predators (Samelius and Alisauskas 2001) or diverting attention away from the nest through distraction displays (Bengtson 1970, Jehl 1973). In this way, leaving the nest to feed is linked to certain costs (e.g., cooling of the eggs, increased risks of nest predation) and benefits (e.g., resource acquisition), all of which must be managed in order to successfully hatch a clutch of eggs (Lecomte et al. 2008, Smith et al. 2012). These trade-offs dictating parental decisions are managed through strategies that can differ among species, most notably for mono-parental species in which the incubating parent is never replaced by the other (Reneerkens et al. 2011, Moreau et al. 2018). Whatever the type of parental care, incubation and its trade-offs remain important in the life history of Arctic-breeding birds.

At a larger scale, breeding success is an essential component of population dynamics. Consequently, the breeding period of most shorebirds is especially critical in the current context where over 44% of shorebird populations are declining (Gratto-Trevor *et al*. 2012). This decline is being observed despite the fact that shorebirds could benefit from climate change in the Arctic (McKinnon et al. 2013, Weiser et al. 2018). Other authors have suggested that a phenological mismatch caused by climate change between shorebirds and their prey, terrestrial arthropods (Tulp and Schekkerman 2008), entails negative impacts on the survival of shorebird chicks (McKinnon et al. 2012, Senner et al. 2017). Although the effects of the responses of arthropods to climate change on shorebirds have been investigated in terms of the timing of their emergence, it is possible that this negative effect could be counteracted by a simultaneous increase in the abundance of arthropods. While food supplementation studies on birds are common in behavioral and population ecology (Boutin 1990, Ruffino et al. 2014), to our knowledge, studies have not yet investigated the effects of increased arthropod abundance on incubating arctic shorebirds. Indeed, if shorebirds are limited by their resources, this boosted food source could benefit shorebirds during a critical time of their life cycle by modifying the balance of trade-offs that parents face during incubation.

Our objective is to determine whether an increase in arthropod abundance can lead to positive effects on the reproductive success of an arctic-nesting shorebird. We hypothesize that increases in arthropod abundance will improve the nesting success of female white-rumped sandpipers, a uniparental incubator, by improving their body condition and/or by allowing them to spend more time incubating (Fig. 1). Improved body condition should allow females to deter nest predators more efficiently (Wallin 1987) and reduce the risks of nest abandonment (Bustnes et al. 2002, Weiser et al. 2017) while spending more time incubating should ensure that eggs develop correctly (Olson et al. 2006, Reneerkens et al. 2011) and reduce detection of the nest by predators (Smith et al. 2012). We predict that (P1) the daily rate of mass loss will be smaller for females having received a resource supplementation and/or that (P2) females having received a resource supplementation will have a lower frequency of incubation recesses and/or that (P3) females having received a resource supplementation will have a lower mean duration of incubation recesses. If P2 or P3 is verified, we predict that (P4) females having received a resource supplementation will spend more time incubating per day. Lastly, if P1 or P4 is verified, we predict that (P5) nesting success of females that received a resource supplementation will generally be better than that of females that did not receive such inputs (Fig. 1). Here we conducted an experiment to test our hypothesis by adding food resources available to nesting sandpipers and measuring 5 response variables, which map directly onto our predictions.

**Figure 1.**
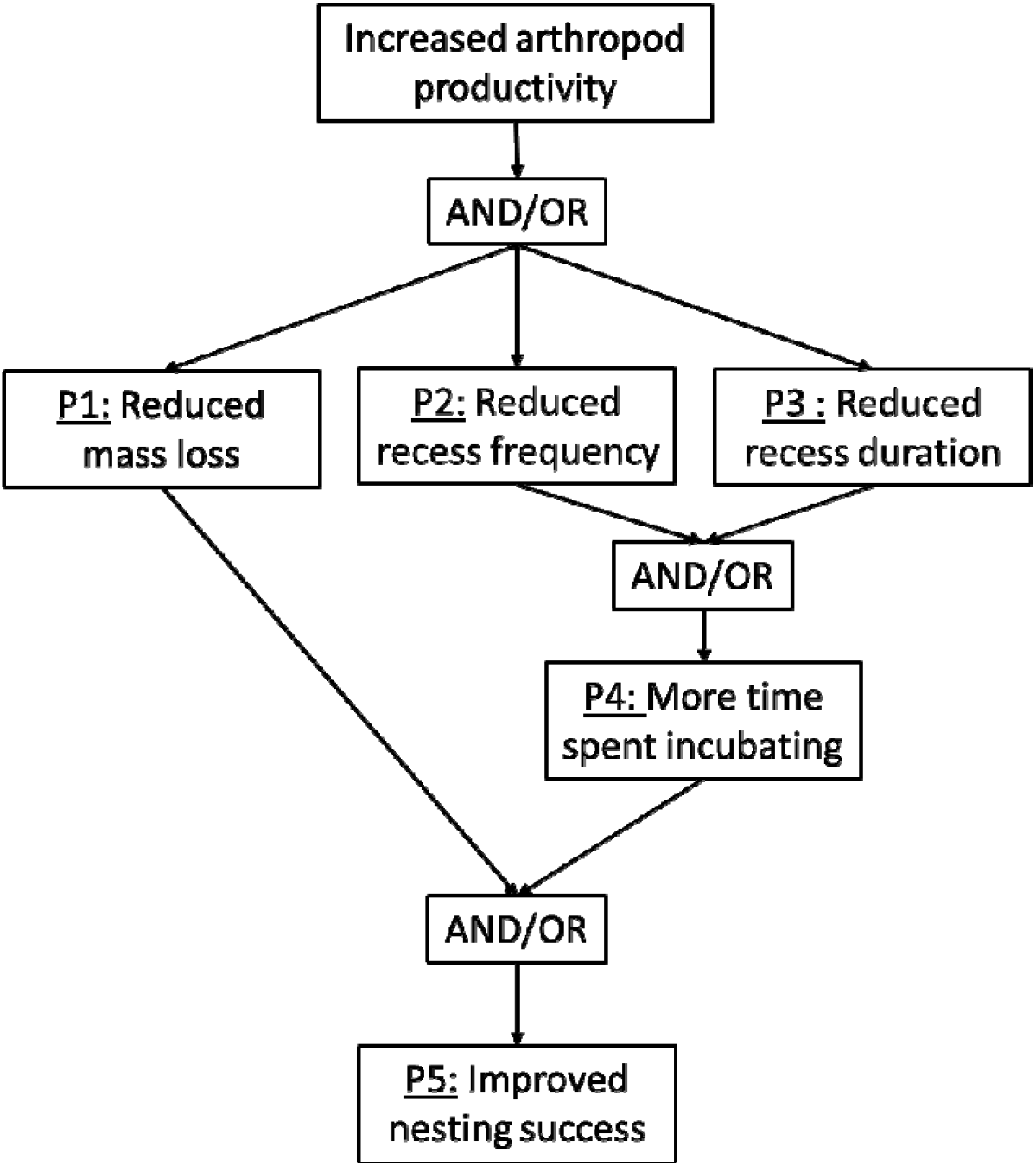
Predictions derived from the hypothesis that increases in arthropod productivity will improve the nesting success of female white-rumped sandpipers by improving their body condition and/or by allowing them to spend more time incubating.

## Materials and methods

### Study species

We chose the white-rumped sandpiper as our model species because they are not globally threatened, they are abundant on our study site, they are income breeders, and they possess typical shorebird morphology and phenology. White-rumped sandpipers are small (15–18 cm, 28–66 g), polygynous, monoparental arctic-nesting shorebirds. Females will lay 4 (occasionally 3) cryptically coloured eggs, and incubation lasts approximately 21 days. As a long-lived and iteroparous species, white-rumped sandpipers may abandon their nests near the end of the breeding season to improve their survival and optimize their southern migration (Weiser et al. 2017).

### Study site

Our experiment was conducted on Igloolik Island (Nunavut, Canada, 69 °24’N, 81 °32’W; Fig. 2a, b), an island in the Foxe Basin, located north-east of the Melville Peninsula during the summers of 2016 and 2017. Our study site covered an area of approximately 25 km^2^ consisting of wet meadows and raised beaches, which comprise mostly xeric and mesic ridges. Wet meadows, interspersed by lakes and ponds, dominate the landscape (Forbes *et al*. 1992) and provide optimal breeding grounds for the two most abundant shorebirds of the island: white-rumped sandpipers (*Calidris fuscicollis*) and red phalaropes (*Phalaropus fulicarius*). Most shorebirds nesting on the island feed on terrestrial spiders and insects. In terms of avian nest predators, shorebirds share the island with nest predators such as long-tailed and parasitic jaegers (*Stercorarius longicaudus*and *Stercorarius parasiticus*), glaucous gulls (*Larus hyperboreus*), Sabine’s gulls (*Xema sabini*), herring gulls (*Larus argentatus*), Thayer’s gulls (*Larus thayeri*), ruddy turnstones (*Arenaria interpres*), and common ravens (*Corvus corax*). Terrestrial predators such as arctic foxes (*Vulpes lagopus*) and stoats (*Mustela ermine*) can also depredate nests. Although these two mammals rarely depredate adults, peregrine falcons (*Falco peregrinus*), gyrfalcons (*Falco rusticolus*), and snowy owls (*Bubo scandiaca*) more frequently prey upon adult shorebirds.

**Figure 2.**
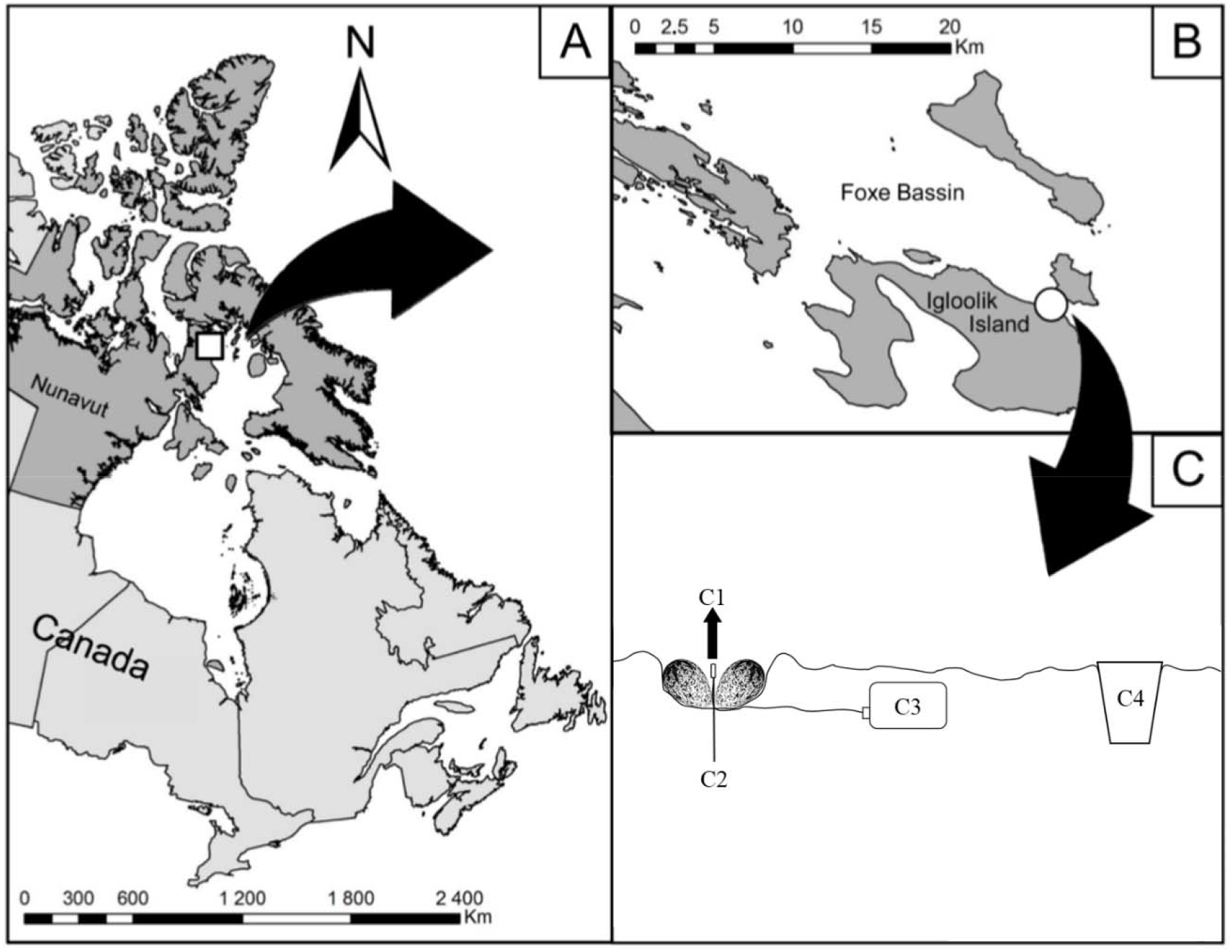
Location of the study area on Igloolik Island (Nunavut, Canada, 69.39 N; 81.55 W). Location of the study area (white circle in [B]) on Igloolik Island (Nunavut, Canada, 69.39 N; 81.55 W); (A and B)). (C) Cross-sectional view of the design of the supplementation experiment conducted during the summers of 2016 and 2017 on and around white-rumped sandpiper nests. We placed a small thermistor (C1) attached to a thin wooden stick (C2) in the middle of the nest. We buried or covered the data logger (C3) and wire. We placed a small cup (full of mealworms or empty) in a hole 5 meters away from the nest.

### Nest monitoring

During the summers of 2016 and 2017, we searched for white-rumped sandpiper nests that were in their laying or incubation stages from around mid-June until our departure at the end of July over an area of 10 km^2^. Nine randomly located circular plots (200m radius) were defined to estimate nest densities although our nest searching was not limited to these plots. When nests were found during the laying stage (nests had more eggs during subsequent visits), we calculated initiation date by considering that one egg was laid per day; hatching date was then estimated to occur 24 days later (3 days of laying and 21 days of incubation). To estimate initiation dates and hatching dates for nests found during the incubation phase, we floated two to three eggs (van Paassen et al. 1984). For nests that eventually hatched, we corrected this date by subtracting 21 days of incubation and 3 days of laying from the hatch date if the latter was available. We also installed a small thermistor (Tinytag Talk PB-5005-0M6 [ø 2.5mm] with a Talk 2 logger TK-4023, Gemini Data Loggers Ltd, Chichester, United Kingdom) within the nest by attaching it to a thin wooden stick planted at the center of the nest (Fig. 2c). The thermistor wire and the data logger were both buried to avoid attracting predators. Prior to deployment, we programmed the logger of each thermistor to start recording temperature every minute for 21 days. Because white-rumped sandpiper females incubate for 21 days (Parmelee, 1992) and because it was unlikely that we started the experiment on the day the clutch was completed, thermistors never stopped recording before the end of incubation. We revisited nests every 4 days and, when approaching the estimated hatching date, every 2 days to be able to track nesting success accurately. We considered that nests in which at least one egg had hatched were successful. Once a nest was found empty, we searched for clues that could indicate whether the nest was successful or not, as well as why the incubation might have failed (Brown *et al*., 2014). Some of these clues include the presence of hatchlings, eggshell fragments or halves, as well as fox urine and scat. We also removed the thermistor and data logger at this time and downloaded the data using the tinytag explorer 4.8 software (Gemini Data Loggers). We cross-checked the fate determined through nest monitoring with the profiles of incubation obtained from the thermistors. Around the midway point of the study period, we visited each nest to estimate vertical cover of vegetation over the nest cup (nest cover; estimated in increments of 10%) and soil humidity (dry, slightly humid, saturated, flooded) over two days.

### Supplementation experiment

The experiment started as soon as possible with the random designation of control or supplementation treatments to nests as they were discovered. We chose to use industrially sold dried mealworms (*Tenebrio molitor*) as our supplementation because they have previously been used as experimental resources for little stints (Tulp and Schekkerman, unpubl. data) and red knots (Vézina, unpubl. data) and are commonly used to feed captive or rescued shorebirds (e.g., Gartell et al. 2014). Additionally, we observed white-rumped sandpipers feeding on mealworms (video available along with data and code on figshare taken during preliminary tests in 2015; see link at the end).

For all supplemented nests (n = 15 in 2016, n = 13 in 2017), we placed a plastic cup (~4 cm diameter) filled mealworms 5 m east of the nest (Fig. 2c). Approximately half of the cup was buried in the ground but remained accessible to sandpipers. This supplementation was available to the sandpipers for the rest of their incubation; since refills were not required required, we considered the number of mealworms as *ad libitum* resources for the breeding individual white-rumped sandpiper. We also used a cup containing only a rock as a weight to maintain it in the ground and set the cup 5 m east of each control nest (n = 14 in 2016, n = 15 in 2017) to account for any positive or negative bias related to the installation of the cup. We later estimated that the average nest age was 8 days (SD = 4.7; n = 57) at the initiation of the experiment.

In 2016, we used bownets to capture females on their nest. We banded each female, evaluated their body condition, and measured morphometric dimensions at the beginning of the experiment. We measured mass using a hanging Pesola scale (±0.5 g precision), the lengths of the head, culmen, and tarsus using a caliper (±0.1 mm), and wing length using a ruler (±1 mm). Females were recaptured and weighed towards the end of the incubation period: on average, 10 days after the first capture. Only in 2017, 11 females were captured as part of other protocols.

### Mass loss models (P1)

We calculated a daily rate of mass loss for each incubating female by subtracting their initial weight from their final weight and dividing the result by the number of days between the two captures. Head, culmen, and tarsus lengths were correlated (Pearson’s r = 0.82, 0.59, and 0.49) and were combined by extracting the principal axis from their PCA to obtain a metric of bird size. In all analyses, nest cover was converted into a categorical variable with two levels: 0-10% and 20-40%. To test for evidence in support of P1, we identified 8 fixed-effects linear models with mass loss as their response variable and which comprised the main predictor (categorical; supplementation and control) and different covariates (Table S1). To limit the complexity and number of models, we defined concurrent models based on categories of covariates. Three models used environmental covariates: soil humidity (categorical; dry and wet), and nest cover (categorical; 0-10% and 20-40%). Two models used biological covariates: nest initiation date (numeric; Julian days), initial weight (numeric; grams), and size (numeric; principal axis of PCA, see above). We expected the supplementation’s effects to be modulated by soil humidity or initiation date, so we included their interaction terms. Finally, two models contained no covariates.

### Incubation models (P2, P3, and P4)

For each dataset of nest temperatures recorded by the thermistors, we computed a moving median and considered that a bird had vacated its nest if temperatures were 3 °C below this threshold. We then calculated the average number of recesses per day per nest over the study period (hereafter recess frequency) by dividing the total number of recesses by the total number of minutes of data and multiplying the result by 1,440 minutes/day. We divided the total number of recess minutes by the total number of recesses to obtain the average duration of recesses (hereafter recess duration). Finally, we calculated the proportion of time spent incubating (hereafter nest attendance) by dividing the total number of incubation minutes by the total number of minutes of data.

We investigated the relationships between recess frequency and recess duration using type 2 linear regressions. Using type 1 linear models, we examined whether nest attendance could be accurately predicted by either or both previous metrics. To test for evidence in support of P2, P3, and P4, we identified three sets of 6 fixed-effects models with recess frequency, recess duration, and nest attendance, respectively, as their response variable (Tables S1-S2-S3, respectively). Aside from the null model, these models comprised combinations of the main predictor (supplementation) with categories of covariates. Three models contained environmental variables, soil humidity (categorical; dry and wet) and nest cover (categorical; 0-10% and 20-40%), and one model contained the biological variable, nest initiation date (numeric; Julian days). We expected the supplementation’s effects to be modulated by soil humidity or initiation date, so we included their interaction terms. Finally, two models contained no covariates.

### Nesting success models (P5)

Nesting success was treated as a binomial variable indicating success (at least one hatched chick) or failure (no hatched chicks) of the incubation. We evaluated nesting success using the RMark package (Laake 2013) to estimate daily survival probabilities over time and compensate for apparent nesting success by considering exposure days (Mayfield 1961, 1975). To test for evidence in support of P5, we identified 10 fixed-effects models (Table S5) of daily nest survival with different combinations of the main predictor (supplementation) and categories of covariates. We included the year variable in all models because we observed a large increase in predator abundance in 2017 accompanied by widespread and frequent nest failures across most shorebird species monitored. We also included the interaction between time and the supplementation in all models because we expected that the effect of the latter would vary during the study period. Four models included combinations of time (numeric; days), year (categorical; 2016 and 2017), and supplementation variables only. A single model included nest cover (categorical; 0-10% and 20-40%) in addition to the previous variables, while four models included biological variables, nest attendance (numeric), recess frequency (numeric), and recess duration (numeric), instead. Only the null model contained no covariates. Due to a lower sample size available for its analysis, we separately investigated the effect of mass loss on nesting success through a fixed effect model of daily nest survival including mass lost per day and time as covariates.

We retained the top-ranked model from each set of models (mass loss, recess frequency, recess duration, nest attendance, and nest survival) using model selection via AIC with second-order bias correction (Burnham and Anderson 2002). We also closely inspected other models that obtained similar AICc scores while checking for potential pretending variables (variables that do not improve the fit of the model and whose estimates overlap 0; Anderson 2008). All statistical analyses were performed with R version 3.2.2 (R Development Core Team 2015), and AlCc model selection was conducted using the AICcmodavg package (Mazerolle, 2017).

## Results

### General results

During the summer of 2016, we found 55 white-rumped sandpiper nests that were in their laying or incubation stages, versus 45 in 2017. Within our 9 randomly located nest searching plots, this equates to 23 and 16 nests/km^2^ in 2016 and 2017, respectively. A total of 57 nests were used in our experiment (29 in 2016 and 28 in 2017), as the others were either part of other projects or were empty before we could set up the experiment. Monitored nests were generally initiated during the final week of June.

### Mass loss analyses (P1)

White-rumped sandpipers weighed 46.0g (SD = 3.5g, CV = 0.08) and 44.8g (SD = 2.7g, CV = 0.06) on average at the beginning and end of the experiment, respectively. Changes in mass were variable during the experiment: while some females lost up to 12.5 g, others gained up to 6.0 g across the studied period. The top-ranked model explaining variation in mass loss per day (R^2^_adjusted_ = 0.44, Table S1) included the supplementation, initial weight, and their interaction (Table 1). We found no support for P1 since the resource supplementation had no impact on the rate of mass loss of incubating females (95% CI = [−0.10, 0.32], Fig. 3a) regardless of initial weight. However, we found that birds lost weight proportionally to their initial weight (95% CI = [0.06, 0.43], Fig. 3b), regardless of whether they received a resource supplementation (95% CI = [−0.24, 0.21]). Heavier birds at the start of the experiment lost more weight per day: for example, our model predicts that a 41g female will gain 0.29g/day while a 51g female will lose 0.41g/day. We also found that an extreme value in our dataset (a 57g female that lost 1g/day) was not a leverage point (Cook 1979) as it did not substantially change the outcome of the analysis.

**Figure 3.**
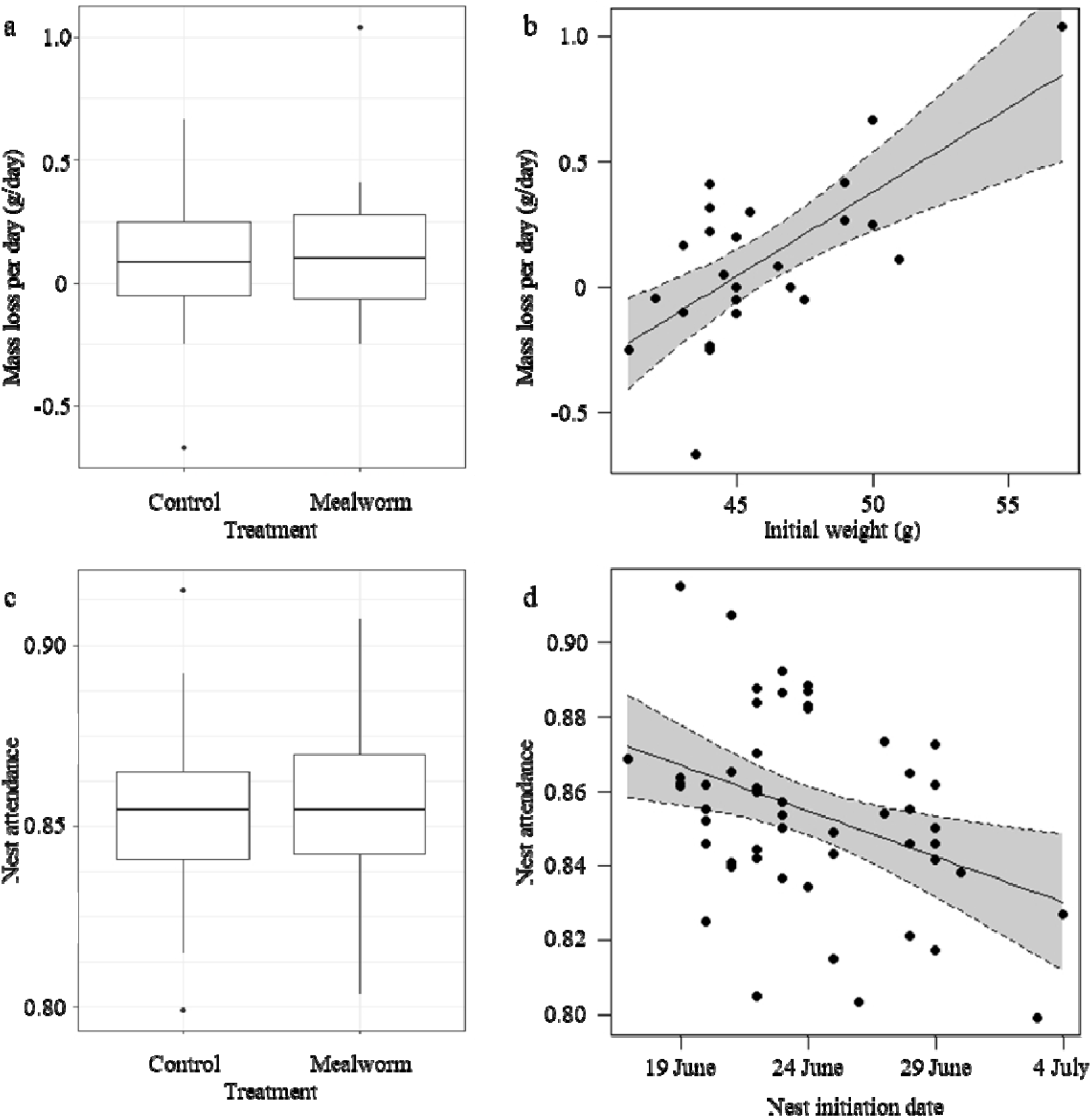
Effect of a resource supplementation (mealworms) and of other covariates on mass loss and nest attendance of incubating white-rumped sandpiper females. Effect of the resource supplementation on mass loss per day (a). Effects of initial weight of incubating females on mass loss per day (b). Effect of resource supplementation on nest attendance (c). Effects of the date at which females started incubating on nest attendance (d). The “whiskers” of the box extend from the hinge to the largest and smallest values at most 1.5 x the inter-quartile range of the hinge. Dotted curves represent 95% CI.

**Table 1.** Parameter estimates from the top-ranked multivariate fixed-effects linear models examining the impact of a resource supplementation on mass loss, nest attendance, and nesting success of incubating white-rumped sandpiper females using the information-theoretical approach.

|  | Estimate | CI (lb) | CI (ub) | SD |
| --- | --- | --- | --- | --- |
| **Mass loss model (P1)** |  |  |  |  |
| Intercept | 0.06 | -0.09 | 0.20 | 0.07 |
| Supplementation | 0.11 | -0.10 | 0.32 | 0.10 |
| Weight | 0.25 | 0.06 | 0.43 | 0.09 |
| Supplementation x Weight | -0.02 | -0.24 | 0.21 | 0.11 |
| **Recess frequency model (P2)** |  |  |  |  |
| Intercept | 27.4 | 25.8 | 29.0 | 0.8 |
| **Recess duration model (P3)** |  |  |  |  |
| Intercept | 7.8 | 7.4 | 8.2 | 0.2 |
| **Nest attendance model (P4)** |  |  |  |  |
| Intercept | 0.858 | 0.848 | 0.867 | 0.005 |
| Supplementation | -0.005 | -0.018 | 0.008 | 0.007 |
| Initiation | -0.013 | -0.023 | -0.003 | 0.005 |
| Supplementation x Initiation | 0.009 | -0.005 | 0.023 | 0.007 |
| **Nesting success model (P5)** |  |  |  |  |
| Intercept | 4.68 | 3.78 | 5.59 | 0.46 |
| Year | -1.83 | -2.75 | -0.91 | 0.47 |
| Time | -1.91 | -2.75 | -1.07 | 0.43 |

Subscripts. Supplementation: experimental addition of resources near the nest; Weight: weight of the incubating female at the beginning of the experiment (standardized); Initiation: Julian date at incubation started (standardized); Year: 2017 compared to 2016 (reference level); Time: number of days since the start of the experiment (standardized); Estimate: predictor's estimate; CI (lb): lower bounds of the confidence interval; CI (ub): upper bounds of the confidence interval; SD: standard deviation.

### Incubation analyses (P2, P3, and P4)

Over the course of our experiment, we found that female white-rumped sandpipers incubated for 85.5%of the time (SD = 2.5). On average, they left their nests 27.4 times per 24 hours (SD = 5.7). Such recesses lasted 7 minutes and 48 seconds on average (SD = 1.4). Among incubating females, nest attendance displayed very little variability (CV = 0.03) compared to recess frequency (CV = 0.21) and duration (CV = 0.18, Fig. S1).

Recess frequency and duration were negatively correlated and shared 37.5% of variability. Unsurprisingly, recess frequency and recess duration together explained 95.1% (F-statistic = 490.8) of the variability in nest attendance. Separately, they only explained respectively 27.4% (F-statistic = 20.35) and 8.2% (F-statistic = 5.5). We found no support for P2 or P3 because the top-ranked models for recess frequency and recess duration were the null models which contained no variables (Tables F-G). The top-ranked model explaining variation in nest attendance (R^2^_adjusted_ = 0.10, Table S4) included treatment, nest initiation date, and their interaction (Table 1). We found no support for P4 since the resource supplementation had no impact on nest attendance of incubating females (95% CI = [−0.018, 0.008], Fig. 3c). However, we found that nest attendance decreased with the initiation date of nests (95% CI = [−0.023, −0.003], Fig. 3d), regardless of whether they received a resource supplementation (95% CI = [−0.005, 0.023]). According to this model, a 10-day difference in the beginning of incubation would result in a 3% difference in the time spent incubating. Extrapolated over a 21-day long incubation period, this would equate to a difference of 15 hours and 7 minutes of incubation.

### Nesting success analyses (P5)

We successfully determined the fate of 50 of the 57 nests used in our experiment. In 2016, 19 nests were successful while 7 failed. In 2017, only 5 nests were successful and 19 failed for a total of 24 successful clutches out of 50 across both years. On average, nests were initiated around June 23^rd^, hatched around July 15^th^, and failed around July 11^th^. Among nests that received the mealworm supplementation, 15 failed and 12 hatched, as compared to 14 failed and 14 hatched control nests. The top-ranked model explaining variation in daily nest survival included time (days since the first nest) and the year of the experiment (Tables 1 and S5). We found no support for P5 since the resource supplementation variable was not included in this model. However, we found that daily nest survival was lower in 2017 compared to 2016 (95% CI = [−2.75, −0.91], Fig. 4a) and decreased over time (95% CI = [−2.75, −1.07], Fig. 4b).

**Figure 4.**
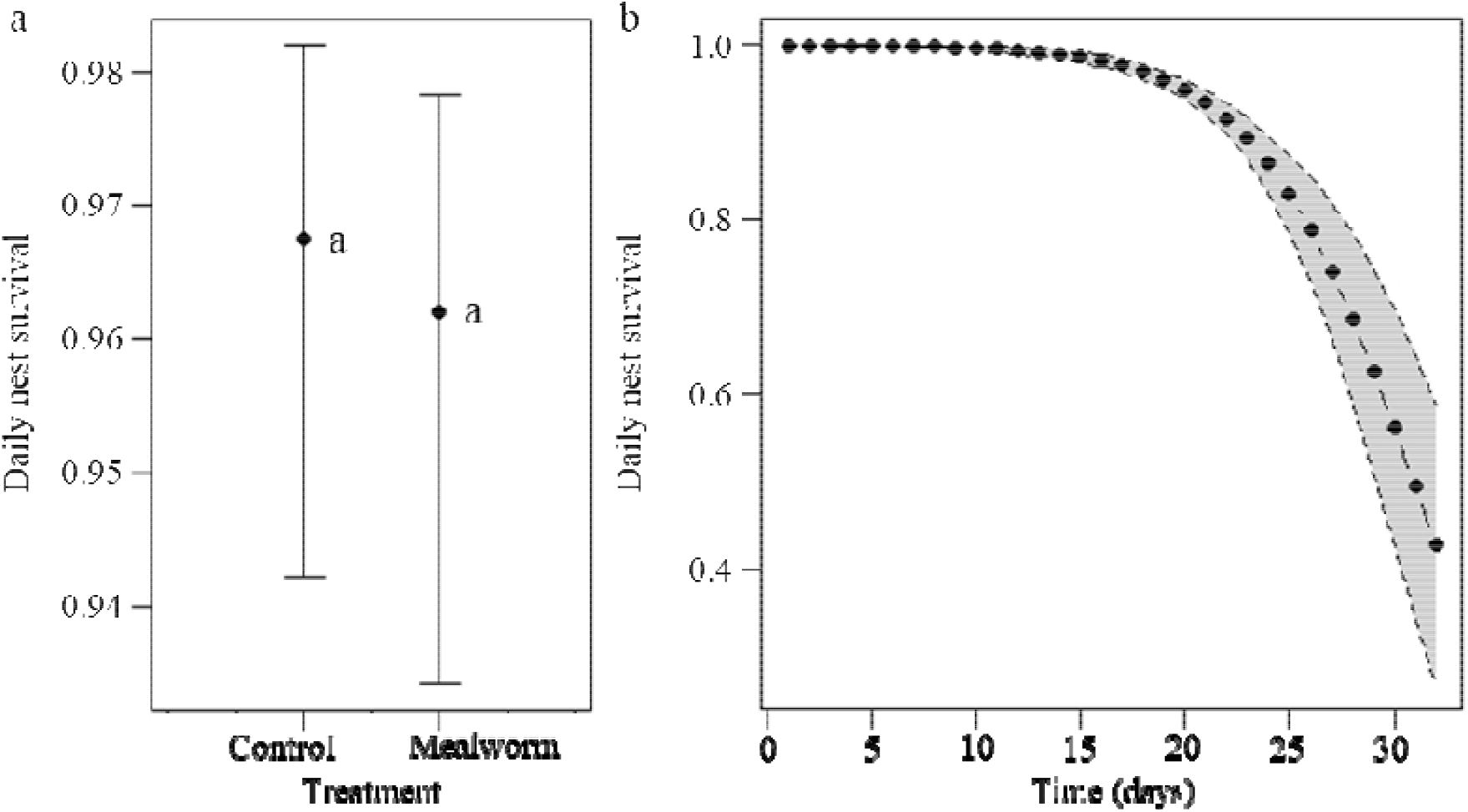
Effects of a resource supplementation (mealworms; a) and time (b) on the daily survival rate of white-rumped sandpiper nests. Groups sharing the same letter are not different according to our top-ranked fixed-effects model (see Table 3). Error bars represent 95% confidence intervals. Grey areas represent the standard error.

## Discussion

We carried out a supplementation experiment mimicking an increased abundance of arthropods in the Canadian Arctic and found no evidence that white-rumped sandpipers should be affected by additional resources while nesting. Contrary to our predictions, the rate of mass loss per day, nest attendance, and nesting success of incubating females were all unaffected by resource supplementation. This does not support our hypothesis that increases in arthropod productivity during incubation could increase the nesting success of female white-rumped sandpipers by improving their body condition and/or by allowing them to spend more time incubating. However, we found that mass loss per day, nest attendance, and nesting success were likely linked to the initial reproductive traits of incubating females. This suggests that *in situ* resources may not be limiting for white-rumped sandpipers during incubation, yet the strategies that they employ during this period, and the successfulness of the latter, seem to be driven by inter-individual traits upon the initiation of incubation. We will explore how these traits impact breeding as well as why our supplementation experiment had no detectable effects in the following sections.

### Mass loss (P1)

In an experiment carried out on the tropical wintering site of a migratory bird, Cooper *et al*. (2015) observed that diminished abundances of arthropod prey had no effect on body mass. However, birds that were exposed to food reduction deposited more fat and lost more pectoral muscle mass, leading to delayed departure dates for spring migration (Cooper et al. 2015). Similarly, it is possible that our supplementation, while leaving mass loss unchanged, had an effect on the body composition of white-rumped sandpipers (Gloutney et al. 1999). Compared to the control group, females that had access to additional resources may have been able to develop to a greater extent their flight muscles in preparation for their fall migration starting only a few days after chicks have hatched. This could then lead to positive carry over effects on adult survival during migration or even wintering. Although authors have addressed the possibility of carry-over effects of resource abundance affecting migrating birds on their breeding grounds (Marra et al. 1998, Robb et al. 2008, Harrison et al. 2011, Legagneux et al. 2012), to our knowledge, no studies have considered effects originating in the Arctic and whose repercussions would be observed during other stages of the life history of birds.

The positive linear relationship between weight loss and weight has already been shown in birds (Monaghan et al. 1989) and suggests that mass loss of smaller birds is more constrained physiologically than that of larger birds. Because weight loss increased with initial weight and was unaffected by the size of birds, it is likely that heavier females, who lost up to 1g/day, had more body reserves and did not need to feed as much. On the other hand, leaner females gained weight (up to 0.7g/day) during the incubation period, suggesting that they prioritized feeding.

### Incubation (P2, P3, and P4)

We observed a strong negative relationship between recess frequency and recess duration, with a similar negative effects of recess frequency and duration on nest attendance; we also measured a high inter-individual variability in recess frequency and duration while nest attendance among individuals remained relatively constant. These analyses reveal that white-rumped sandpipers can have variable strategies to maintain the balance between the frequency of recesses and their duration. However, because the increase of frequency was generally accompanied by a reduction in duration (and vice versa), the resulting outcome, total nest attendance, varied very little among different individuals (CV = 0.03). In contrast, recess duration and frequency showed larger variability (CV = 0.18 and 0.21 respectively), yet we could not detect any major biological drivers (including the supplementation) of this variability. Because we found no evidence of an effect of the resource supplementation on nest attendance, we suggest that the latter is not dependent on the abundance of resources during the incubation period. Individual characteristics and various predation pressures could be at play and warrant further investigations.

Despite its low variability, nest attendance decreased when delays in nest initiation increased (−0.3% per day). Because early nest initiation is associated with high body condition (Blums et al. 2002, 2005), nest attendance may in fact be driven by the physical state of white-rumped sandpipers at the beginning of incubation. Females that initiated early likely spent more time incubating at the expense of feeding time; their better body condition allowing them to use previously accumulated body stores. Females that initiated later in the season would have spent less time incubating and more time feeding to compensate for their poorer body condition. As it was the case for mass loss, we found evidence that the trade-off between incubation and feeding is unaffected by supplementation, suggesting that this trade-off is rather driven by a parameter linked to the body condition of females at the beginning of incubation.

### Nesting success (P5)

Although recess frequency and nest attendance can partially predict nesting success in shorebirds (e.g. Smith *et al*. 2012, Meyer et al. 2020), we found no evidence of these relationships in our experiment. Declines in nesting success have been linked with increased predation pressure in temperate systems (Patnode and White 1992, Sloan et al. 1998). In the Arctic, nest survival has been suggested to decline due to increasing risks of nest abandonment while predation pressure remains stable throughout the summer (Weiser et al. 2017). Generally, long-lived species favour their survival and may abandon the nest when breeding costs reach critical energy demands (Williams 1966, Weiser et al. 2017). Our results suggest that nest abandonment became more prevalent as the season progressed even though sandpipers had access to resource supplements. This implies that the quantity of resources during incubation is unlikely a signal to leave the nest before hatching. However, shorebirds face unpredictable abiotic (Tulp and Schekkerman 2006, Smith and Wilson 2010) and biotic (Smith et al. 2012) conditions that may raise incubation costs beyond their limit, forcing them to abandon. The fact that our supplementation had no effect on nest survival indicates that white-rumped sandpipers cannot compensate for these unexpected incubation costs with increased nutritional intake. For the future scenarios of increased productivity in the Arctic, we suggest that an increased peak of arthropods during incubation may not necessarily translate into higher nesting success of these sandpipers. However, they could benefit before (increased fueling) and after (prior to migration, chick survival) incubation.

By noting that nest survival declines as the short Arctic summer progresses, we add support to other studies that early initiation favors higher nest survival in birds (Blums et al. 2005, Weiser et al. 2017). As early initiation is often displayed by the birds possessing the best body condition (Blums *et al*. 2002, 2005), we can once again highlight the positive relationship between the state of females at the beginning of incubation and nesting success. Body conditions at the start of incubation can be determined in part during the pre-laying stage when shorebirds reconstitute their body stores post-migration (Drent & Daan 1980; Morrison & Hobson 2004). The pre-laying stage is then likely more critical than the incubation stage in terms of energy gains and repercussions on incubation. We can thus expect additional resources to affect shorebirds during this period. Considering that the decision of whether or not to breed often relies on reaching a threshold of body condition (Rowe et al. 1994, Legagneux et al. 2016), additional resources on the breeding grounds could increase the number of individuals whose body condition reaches such theoretical threshold (Legagneux et al. 2012). In turn, more individuals could invest in breeding (Prevedello et al. 2013) rather than skip reproduction (Legagneux *et al*. 2016).

### Conclusion

We found no evidence of an effect of our resource supplementation on mass loss, incubation, and nesting success of female white-rumped sandpipers during their incubation period in the Canadian Arctic. It is important to highlight that those methodological limitations of this study may contribute to explaining the lack of detected effects. It is possible that the number of mealworms consumed varied from one female to another, reducing our ability to draw conclusions on the effects of the supplementation. While we attempted to quantify the number of mealworms consumed, disturbances such as rain, wind, and flooding made this very difficult. It is also frequent for incubating sandpipers to fly up to 200m away from their nest to feed. With this in mind, it is possible that females only fed upon the supplementary resources when walking to and from their nest, even though the food supplement was always available *ad libitum*.

Other authors have suggested that supplementation experiments tend to have insignificant effects on individuals when they naturally have access to good or moderate amounts of resources (Boutin 1990, Nager et al. 1997, Ruffino et al. 2014). Although this would be difficult to quantify, it is possible that the abundance of arthropods in the Arctic may be above the saturation point of shorebirds during part of the summer. Studies that quantify arthropod abundance systematically as a resource for shorebirds (Schekkerman et al. 2003) could eventually verify whether arctic-breeding shorebirds have access to sufficient, non-limiting resources during incubation. Also, it seems that predation, rather than resource availability, is the leading cause of nest failure in shorebirds (Weiser et al. 2017). The fact that our supplementation experiment had no effects on white-rumped sandpiper females during their incubation adds to the evidence that *in situ* resources are not the principal limiting factor for animals living at high latitudes (Schoech and Hahn 2007, Ruffino et al. 2014). This would limit how the warming of the Arctic could indirectly benefit shorebirds during their incubation through increased abundances of arthropods. Further investigation of the carry-over effects of increased resources in the Arctic on shorebirds will help determine whether some declining populations can find relief in climate warming.

## ETHICS

Our work was approved by the Animal Care Committee of the Université of Moncton.

## DATA ACCESSIBILITY

The data, the video, and the code used in this study are available at https://figshare.com/s/6af0f26709ee8a4b4134.

## AUTHORS’ CONTRIBUTIONS

LM: data curation, formal analysis, writing—original draft, writing—review and editing; NL: conceptualization, funding acquisition, project administration, supervision, analysis, writing—review and editing.

## Acknowledgements

We thank the entire Igloolik field crews from 2016 and 2017 for assistance in finding nests, capturing birds, and collecting data. We thank Université de Moncton, Indian and Northern Affairs Canada, Polar Continental Shelf Project, the Government of Nunavut for logistical support. We thank O. Gilg for helping to secure a good rebate on the Tiny Tags purchase. This work benefited from comments by M.-A. Giroux, S. Leroux, G. Moreau, J. Bêty, É. Gagnon, D. Hamilton, S. Jacques, and L. Carter.

