## Supplementary Tables and Figures for "Free meals during breeding: increased resource access does not benefit Arctic-nesting shorebirds"

### SUPPORTING INFORMATION

Additional Supporting Information is linked to this study:

**Table S1.** Multivariate fixed-effects linear model selection examining the impact of a resource supplementation on mass loss of white-rumped sandpiper females during incubation.

**Table S2.** Multivariate fixed-effects linear model selection examining the impact of a resource supplementation on the frequency of recesses of white-rumped sandpiper females during incubation.

**Table S3.** Multivariate fixed-effects linear model selection examining the impact of a resource supplementation on the duration of recesses of white-rumped sandpiper females during incubation.

**Table S4.** Multivariate fixed-effects linear model selection examining the impact of a resource supplementation on the nest attendance of white-rumped sandpiper females during incubation.

**Table S5.** Multivariate fixed-effects linear model selection examining the impact of a resource supplementation on the daily survival rate of white-rumped sandpiper nests.

**Figure S1.** Distribution of normalized incubation variables (divided by their respective means) of female white-rumped sandpipers. Recess frequency (a) and recess duration (b) are much more variable among females than nest attendance (c).

**Table S1.**

Multivariate fixed-effects linear model selection examining the impact of a resource supplementation on mass loss of white-rumped sandpiper females during incubation using the information-theoretical approach.

| Predictors of mass loss | LogLik | ΔAICc | $R^2_{adj}$ |
|---|---|---|---|
| Supplementation x Weight | 1.4 | 0.0 | 0.44 |
| Supplementation + Cover | -4.2 | 8.1 | 0.17 |
| Null model | -7.6 | 9.4 | 0.00 |
| Supplementation | -7.3 | 11.5 | -0.02 |
| Supplementation x Soil + Cover | -2.7 | 11.8 | 0.19 |
| Supplementation + Size | -5.6 | 14.1 | 0.02 |
| Supplementation x Soil | -7.0 | 16.9 | -0.09 |
| Supplementation x Initiation | -7.1 | 17.1 | -0.10 |

Subscripts. Supplementation: experimental addition of resources near the nest; Weight: weight of the female at the beginning of the experiment; Cover: vertical cover of vegetation over the nest cup; Soil: soil humidity; Initiation: date at which incubation started; LogLik: log likelihood; ΔAICc: delta AIC with second-order bias correction; $R^2_{adj}$: adjusted R-squared.

**Table S2.**

Multivariate fixed-effects linear model selection examining the impact of a resource supplementation on the frequency of recesses of white-rumped sandpiper females during incubation using the information-theoretical approach.

| Predictors of recess frequency | LogLik | ΔAICc | $R^2_{adj}$ |
|---|---|---|---|
| Null model | -164.1 | 0.0 | 0.00 |
| Supplementation | -164.1 | 2.1 | -0.02 |
| Supplementation + Cover | -164.0 | 4.2 | -0.03 |
| Supplementation x Soil | -163.0 | 4.8 | -0.04 |
| Supplementation x Initiation | -163.5 | 5.8 | -0.02 |
| Supplementation x Soil + Cover | -163.5 | 8.3 | -0.06 |

Subscripts. Supplementation: experimental addition of resources near the nest; Cover: vertical cover of vegetation over the nest cup; Soil: soil humidity; Initiation: date at which incubation started; LogLik: log likelihood; ΔAICc: delta AIC with second-order bias correction; $R^2_{adj}$: adjusted R-squared.

**Table S3.**

Multivariate fixed-effects linear model selection examining the impact of a resource supplementation on the duration of recesses of white-rumped sandpiper females during incubation using the information-theoretical approach.

| Predictors of recess duration | LogLik | ΔAICc | $R^2_{adj}$ |
|---|---|---|---|
| Null model | -89.8 | 0.0 | 0.00 |
| Supplementation | -89.8 | 2.2 | -0.02 |
| Supplementation + Cover | -89.8 | 4.5 | -0.04 |
| Supplementation x Soil | -88.8 | 5.1 | -0.06 |
| Supplementation x Initiation | -89.8 | 7.0 | -0.02 |
| Supplementation x Soil + Cover | -89.8 | 9.5 | -0.08 |

Subscripts. Supplementation: experimental addition of resources near the nest; Cover: vertical cover of vegetation over the nest cup; Soil: soil humidity; Initiation: date at which incubation started; LogLik: log likelihood; ΔAICc: delta AIC with second-order bias correction; $R^2_{adj}$: adjusted R-squared.

**Table S4.**

Multivariate fixed-effects linear model selection examining the impact of a resource supplementation on the nest attendance of white-rumped sandpiper females during incubation using the information-theoretical approach.

| Predictors of nest attendance | LogLik | ΔAICc | $R^2_{adj}$ |
|---|---|---|---|
| Supplementation x Initiation | 122.3 | 0.0 | 0.10 |
| Null model | 118.1 | 1.4 | 0.00 |
| Supplementation | 118.2 | 3.4 | -0.02 |
| Supplementation + Cover | 118.8 | 4.6 | -0.01 |
| Supplementation x Soil | 119.2 | 6.2 | -0.02 |
| Supplementation x Soil + Cover | 119.5 | 8.3 | -0.03 |

Subscripts. Supplementation: experimental addition of resources near the nest; Initiation: date at which incubation started; Cover: vertical cover of vegetation over the nest cup; Soil: soil humidity; LogLik: log likelihood; ΔAICc: delta AIC with second-order bias correction; $R^2_{adj}$: adjusted R-squared.

**Table S5.**

Multivariate fixed-effects linear model selection examining the impact of resource supplementation on the daily survival rate of white-rumped sandpiper nests using the information-theoretical approach.

| Predictors of daily survival | Loglik | ΔAICc |
|---|---|---|
| Year + Time | 184.0 | 0.0 |
| Year + Supplementation x Time + Incubation | 178.1 | 0.2 |
| Year + Supplementation x Time + RecessFreq | 179.3 | 1.4 |
| Year + Supplementation * Time + RecessDur + RecessFreq | 177.7 | 1.8 |
| Year + Supplementation x Time | 181.8 | 1.9 |
| Year + Supplementation x Time + Cover | 180.1 | 2.2 |
| Year + Supplementation x Time + RecessDur | 181.8 | 3.9 |
| Year | 206.7 | 20.8 |
| Year + Supplementation | 206.2 | 22.2 |
| Null model | 222.6 | 34.6 |

Subscripts. Year: year of the experiment; Time: days since the first nest visit of the first nest found; Supplementation: experimental addition of resources near the nest; Incubation: proportion of time spent incubating; RecessFreq: daily frequency of incubation recesses; RecessDur: mean duration of incubation recesses; Cover: vertical cover of vegetation over the nest cup; LogLik: log likelihood; ΔAICc: delta AIC with second-order bias correction.

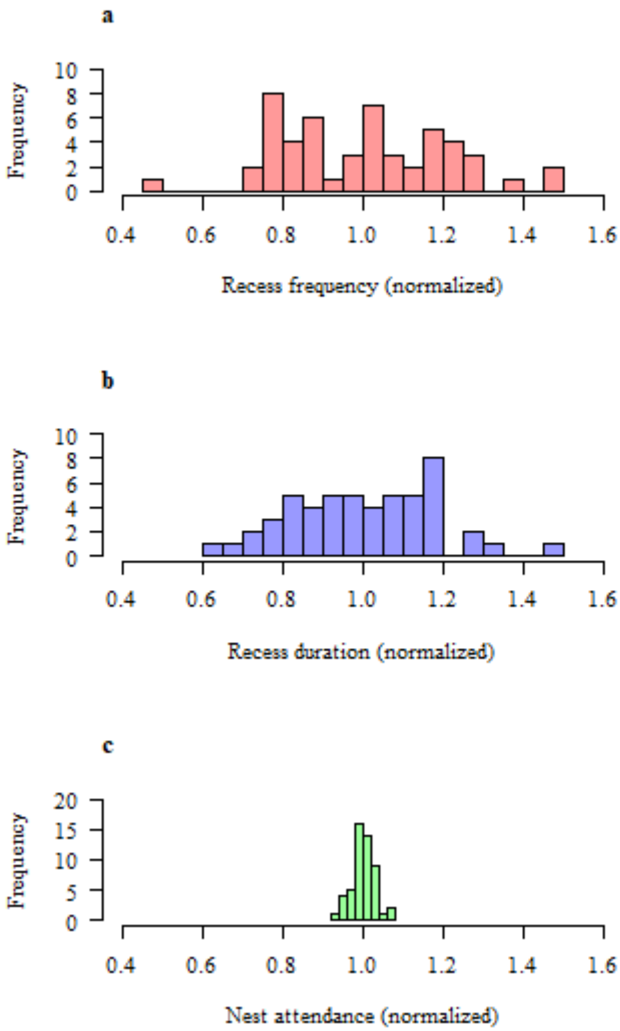


**Figure S1.** Distribution of normalized incubation variables (divided by their respective means) of female white-rumped sandpipers. Recess frequency (a) and recess duration (b) are much more variable among females than nest attendance (c).
